# Effect of pH on the secretome profile of the human pathogen *Candidozyma auris*

**DOI:** 10.64898/2026.04.14.718400

**Authors:** Asier Ramos-Pardo, Guillermo Quindós, Elena Eraso, Elena Sevillano, Vladimir R. Kaberdin

## Abstract

Secreted virulence factors (e.g., hydrolytic enzymes, toxins, agglutinins) play an important role in human diseases. Nevertheless, their secretion by some pathogenic fungi, especially some virulent *Candida-*related species such as *Candidozyma auris,* is still only partly characterized.

Here we used high-throughput mass-spectroscopy analysis to identify polypeptides secreted by *C. auris* into growth medium under two physiologically relevant pH conditions: pH 5.5 and pH 7.5. This analysis revealed that many secreted polypeptides belong to putative virulence factors and enzymes involved in cell wall biogenesis. Moreover, we found that 13 and 27 polypeptides were detected only at pH 5.5 or pH 7.5, respectively. Furthermore, our findings indicate that lower pH (pH 5.5) favours secretion of several putative virulence factors including aspartic proteases and polypeptides potentially facilitating host-pathogen interactions. In contrast, the majority of polypeptides detected only at pH 7.5 are involved in N-glycosylation and protein folding.

Thus, this secretome analysis reveals numerous *C. auris* polypeptides with putative roles in infection and host-pathogen interactions. Moreover, their differential secretion at pH 5.5 and pH 7.5 may reflect different strategies used by *C. auris* to elicit infections in different anatomical sites.

## 1. Introduction

*Candidozyma auris* (formerly *Candida auris*) is a pathogenic yeast that causes severe infections associated with high mortality rates [1]. It has recently led to numerous hospital outbreaks worldwide, which have been very difficult to control [1]. Recent studies demonstrate that the treatment of *C. auris* infections is complicated by the ability of this pathogen to (i) persist in the environment for long time, (ii) show resistance to commonly used disinfectants and (iii) easily survive at elevated temperatures (for a review, see [2]), thereby being a potentially constant source of infection in the healthcare settings (reviewed in [3]). One of the most concerning aspects of *C. auris* is its intrinsic resistance to multiple antifungal drugs, including fluconazole (87-100%), amphotericin B (8-35%), and echinocandins (0-8%). The notorious multidrug resistance along with the challenges in accurate identification of this pathogen through conventional laboratory techniques further complicate timely diagnosis and leave very limited options for effective management of *C. auris* infections [4]. Owing to high threat to human health and few treatment options, the World Health Organization has recently included *C. auris* in the critical priority group of the fungal pathogens [5].

Recent studies have shown that *C. auris* can cause infections in diverse body sites, especially in immunocompromised or high-risk individuals. The initial exposure to this pathogen occurs via direct person-to-person contacts or through contaminated surfaces, subsequently leading to host colonization within days to weeks [6]. The severity and symptoms vary according to the infection site with bloodstream infections representing the most critical cases of *C. auris-*associated candidiasis [7,8]. Unlike closely related *Candida* species normally present in the microbiota of the gastrointestinal tract, *C. auris* uses skin as its most common reservoir, being capable of prolonged persistence on human skin under different acidic conditions [7,9]. Regarding the pathogenic properties, *C. auris* demonstrates a unique resilience compared to *Candida albicans*, which is efficiently recognized and eliminated by the human immune system. This resilience makes *C. auris* a significant threat during bloodstream infections, where it can evade immune defences and cause severe candidemia [10].

A number of *C. auris* virulence factors are believed to greatly affect the infection process by not only enhancing the adhesion capacity of this pathogen but also by facilitating biofilm development and evasion of the host immune response. For instance, it has been shown that *C. auris* can produce the SCF1 adhesin, which forms exceptionally tight contacts, thus enhancing the pathogen’s ability to adhere to host tissues and medical surfaces [11]. Among the *C. auris* arsenal of virulence factors [3], hydrolytic enzymes play a particularly important role in host-pathogen interactions. The vast majority of these enzymes are represented by proteases and lipases [12]. Among secreted *Candida* enzymes with proteolytic activities, aspartic proteases (SAPs) are particularly notable [13]. In *C. albicans*, they are encoded by *SAP* genes (i.e., *SAP1* to *SAP7* genes), whose expression yield aspartic proteases with distinct biochemical characteristics [14] and biological functions [13]. Recent studies have demonstrated that some aspartic proteases also play an essential role in *C. auris* due to their contribution to disruption of the host immune system by degrading host factors (e.g., IgG heavy chains and C3 protein). Notably, the recent study by Kim *et al.* [15] highlights Sap3 as the most significant aspartic protease of *C. auris*. The authors show that this enzyme not only weaken the host immune defences but also undermines the structural integrity of host tissues [15].

Lipases belong to another class of important hydrolytic enzymes involved in both degradation and synthesis of various lipids [16]. Moreover, phospholipases play a particularly prominent role among these enzymes [17]. They contribute to host-pathogen interactions by facilitating the pathogen’s adhesion as well as evasion of the host immune system [18–20]. Although *C. auris* lipases are less studied than those of other *Candida*-related species, several lipases homologous to those of *C. albicans* (e.g., Lip1 and a homologue of CaLip2 [19]) have also been identified in *C. auris.* In addition to proteases and lipases, there are other relevant hydrolases such as glucosidases, which play an essential role in conferring cell wall integrity and plasticity [21]. These enzymes have ancillary functions, being primarily involved in modifications of the pathogen’s cell wall. Similar to *C. albicans*, *C. auris* also produces a number of glucosidases, among which the most abundant ones are the 1,3-β-glucosidase Xog1 and α-1,2-mannosyltransferase MN21 [21]. According to recent studies, *C. albicans* glycosidases appear to be involved in removing certain pathogen-associated molecular patterns (PAMPs) from its own cell wall [22,23], thus enabling the pathogen to evade detection by the host immune system [24]. A similar mechanism may also occur in *C. auris*, potentially contributing to its ability to evade host defences.

Although a number of secreted polypeptides have been classified in *C. albicans* and in some other *Candida* species as essential virulence factors, many their counterparts secreted by a related pathogen, *C. auris*, to elicit infection and to develop diseases remain largely unknown. To provide a wider overview of the secreted polypeptides potentially acting as putative virulence factors as well as to reveal alterations in their production depending on the physiological pH of the typical sites of infection (i.e., skin (pH 4.7-5.7) [25] and blood (pH 7.35-7.45 [26]), we aimed at identifying the *C. auris* polypeptides secreted at two physiologically relevant pH values, namely at pH 5.5 and 7.5.

## 2. Materials and methods

### 2.1. Culturing and storage of C. auris isolates

In this study, we analysed the secretion of hydrolytic enzymes using five clinical isolates of *C. auris* (17-257, 17-259, 19-060, 19-061, 19-063) obtained from La Fe Hospital (Valencia, Spain). All isolates belong to the *C. auris* clade III [27]. Positive controls for protease secretion included *C. albicans* NCPF 3153 and *C. tropicalis* NCPF 3111 (National Collection of Pathogenic Fungi), while *Candida parapsilosis* ATCC 22019 (American Type Culture Collection) served as the positive control for lipase activity. They were retrieved from the strain repository maintained by the Department of Immunology, Microbiology, and Parasitology of the University of the Basque Country (UPV/EHU). For short-term storage, each isolate was maintained in sterile distilled water at room temperature. For long-term preservation, they were inoculated in tubes containing RPMI medium (Sigma-Aldrich, USA) at pH 7.0, mixed with an equal volume of 80 % glycerol and stored at -80 °C.

### 2.2. Detection of protease and lipase activities by plate assays

#### 2.2.1. Inocula preparation

Fresh *Candida* colonies from Sabouraud Dextrose Agar (SDA) plates were resuspended right before inoculation in sterile 0.85 % NaCl solution (bioMérieux S.A., France) to obtain a turbidity of 0.8 McFarland units (Ramos-Pardo et al., 2023), which corresponds to an approximate concentration of 10^7^ CFU/mL.

#### 2.2.2. Preparation of solid media and inoculation

The medium for protease detection using plate assays contained bacteriological agar (20 g/L), yeast extract (0.1 g/L), yeast carbon base (11.7 g/L) and was supplemented with skim milk (2 g/L). The solid medium for detection of lipases contained malt agar (45 g/L), NaCl (58.4 g/L), CaCl_2_ (0.56 g/L) and Tween 80 (20 mL/L). The pH was adjusted to ensure that the final pH after autoclaving would be 5.5 and 7.5. The medium was then autoclaved at 121 °C for 15 minutes before casting the plates. After homogenizing the fresh inocula (see 2.2.1), 3 and 10 µL of each cell suspension were inoculated onto the agar plates for detection of lipases or proteases, respectively, ensuring that the droplets were sufficiently spaced. The plates were then incubated at 37 °C under aerobic conditions for 6 days (Tween 80-containing plates) or 7 days (skim milk-containing plates). To ensure reproducibility and minimize interplate variability, three independent replicates of each experiment were obtained.

### 2.3. Secretome analysis by mass spectrometry

#### 2.3.1. Secretome preparation

To obtain *C. auris* secretomes, cells were grown in minimal liquid medium containing yeast nitrogen base without amino acids (6.7 g/L) and glucose (50 g/L), as described in previous studies [28,29]. Additionally, to maintain the constant pH throughout the entire process, two different buffer systems were used. Namely, the pH of each medium was maintained by adding 100 mM MES (pH 5.5) or HEPES (pH 7.5). Both MES and HEPES are buffering agents that are biologically neutral [30]. Although these reagents belong to zwitterionic compounds with aminosulfonate moieties (i.e., they are chemically similar), their capacity to maintain pH is limited by two different pH ranges, thus justifying the specific use of MES and HEPES at pH 5.5 and 7.5, respectively.

To prepare the *C. auris* 17-257 secretomes for mass spectrometry analysis, fresh inocula (see 2.2.1) were incubated for 24 h at 37 °C, with continuous shaking at 120 rpm. Then, the culture was transferred to 50 mL Falcon tubes, cooled down on ice, centrifuged at 4000 *g* for 10 min at 4° C and filtered through a 0.22 µm filter. The resulting filtrates were then transferred to Amicon Ultra-15 centrifugal filters (Merck Milipore, USA) with 30 kDa cutoff and centrifuged four times at 4000 *g* for 20 min at 4 °C, discarding the filtrates after each centrifugation. Finally, the residual solutions containing the concentrated secretome samples were individually mixed by pipetting and transferred to 1.5 mL tubes prior to their subsequent analysis by mass spectrometry. The final protein concentration of the samples at pH 5.5 ranged from 0.09 to 0.356 mg/mL, while the concentration of those at pH 7.5 varied between 0.12-to-0.52 mg/mL.

#### 2.3.2. Mass spectrometry analysis

The mass spectrometry (MS) analysis was carried out at the Advanced Research Facilities (SGIker) of the University of the Basque Country (UPV/EHU). Three biological replicates were obtained for each pH (i.e., pH 5.5 and 7.5), thus resulting in six samples in total. Prior to MS analysis, equal amounts of each protein sample were individually precipitated with the 2-D Clean-Up Kit (Cytiva, USA) following the manufacturer’s instructions. The precipitated samples were resolved by SDS-PAGE (1 cm run only), the proteins were visualized by Coomassie blue staining and the gel slices corresponding to the entire lanes were manually excised. The excised gel slices were subjected to in-gel endopeptidase digestion as previously described [31] with minor modifications. Namely, the polypeptides present in gel slices were reduced with dithiothreitol (DTT), alkylated with iodoacetamide and incubated in 50 mM NH_4_HCO_3_ with proteomics grade trypsin (Roche Diagnostics, Switzerland) used at 12.5 ng/μL. Following incubation at 37 °C overnight, the recovered peptides were desalted using C18 Spin Columns (Thermo Fisher Scientific, USA), lyophilized in a Speed-Vac (Thermo Fisher Scientific) and stored at -20 °C. Mass spectrometric analyses were performed on an EASY-nLC 1200 liquid chromatography system interfaced with an Exploris 480 mass spectrometer (Thermo Fisher Scientific) via a nanospray flex ion source. Briefly, desalted peptides were loaded onto an Acclaim PepMap100 precolumn (75 μm x 2 cm, Thermo Fisher Scientific) connected to an Acclaim PepMap RSLC (75 μm x 25 cm, Thermo Fisher Scientific) analytical column. The gradient was created by mixing Buffer A (0.1 % formic acid) and buffer B (0.1 % formic acid in 80 % acetonitrile). Peptides were eluted using the following gradient at a flow rate of 300 nL·min^−1^: 0 min: 100% A, 0 % B; 40 min: 81% A, 19 % B; 53 min: 69% A, 31 % B; 61 min: 59% A, 41 % B; 63 min: 5% A, 95 % B and 73 min: 100 % B. Exploris 480 was operated in a positive ion mode. Survey scans were acquired from 375 to 1500 m/z with a resolution of 60,000 and fragmentation spectra with a resolution of 15,000. Precursor ions were fragmented by higher energy C-trap dissociation (HCD) with a collision energy of 30 %. The maximum injection time was set to 50 ms for survey and Auto for the MS/MS scans, whereas the AGC target values of 300 % and 100 % were used for survey and MS/MS scans, respectively. Singly charged ions, ions with more than 5 charges and ions with unassigned charge state were excluded from MS/MS. Data were acquired using Xcalibur software (Thermo Fisher Scientific) and deposited to the ProteomeXchange Consortium via the PRIDE [32] partner repository with the dataset identifier PXD060071 and doi. 10.6019/PXD060071.

#### 2.3.3. Protein identification and quantification

The acquired raw data files were processed with the MaxQuant [33] software (version 2.1.0.0). The processed MS data were searched against the UniProtKB *C. auris* database (release 2024_04) using the parameters detailed below. Namely, carbamidomethylation of Cys was set as fixed modification, whereas Met oxidation and protein N-terminal acetylation were defined as variable ones. Mass tolerance was set to 8 and 20 ppm at the MS and MS/MS modes, respectively. Enzyme specificity was set to trypsin with a maximum of two missed cleavages. A match between runs was enabled and the false discovery rate for peptides and proteins was set to 1 %. Normalized spectral protein label-free quantification (LFQ) intensities were calculated using the MaxLFQ algorithm. MaxQuant output data were analysed with the assistance of Perseus [34] platform (version 2.0.7.0). Initially, the proteins that were only identified by site, contaminants and reverse hits were removed. The search in the UniProtKB *C. auris* database employing the above parameters, revealed 956 protein groups, each corresponding to a single protein represented in the group by its homologues present in different *C. auris* strains. The resulting protein groups (956 in total) were further examined to remove contaminants (e.g., various keratins and trypsin) as well as reverse proteins and other putative false positives including modified peptides. As a result, 887 protein groups remained, of which only the proteins present in 3 biological replicates of at least one pH condition (688 in total) were selected for further analysis.

For quantification of the identified polypeptides, Label-Free Quantification (LFQ) intensity values were used. These values are normalized intensity profiles generated by the MaxLFQ algorithm within MaxQuant software. The LFQ intensity values were log_2_ transformed and a permutation-based False Discovery Rate (FDR) correction was applied using Perseus software. The analysis employed 250 randomizations using an FDR threshold of 0.01, and an s0 factor of 0.5.

#### 2.3.4. Bioinformatic analysis

Analysis of protein groups using SignalP 6.0 (https://services.healthtech.dtu.dk/services/SignalP-6.0/) revealed that out of 688 protein groups, 123 groups contained signal polypeptides and therefore these proteins were considered as members of the putative *C. auris* secretome. Finally, as proteins within each group belong to different *C. auris* strains and their sequence was nearly identical within each group (i.e., sequence similarity was > 95%), only one representative polypeptide belonging to the strain *C. auris* B8441 (well-annotated in *Candida* Genome and STRING databases; see http://www.candidagenome.org/ and https://string-db.org/, respectively) was included in the final list (Table S1).

## 3. Results

### 3.1. Detection of hydrolytic enzymes secreted by C. auris using plate assays

Plate-based assays to detect secreted protease and lipase activities were carried out at two different pHs (i.e., pH 5.5 and pH 7.5), and the results are summarized in Table 1. We found that protease activity was observed only at pH 5.5 (Table 1). In contrast, lipase activity was detected at both pH conditions tested, thus indicating that *C. auris* can efficiently hydrolyse lipids in a wider range of pH.

**Table 1.** *C. auris* clade III isolates analysed in this study.

| Isolate ID<br>(UPV/EHU) | Lipase activity |  | Protease activity |  | Mass-spec<br>analysis |
| --- | --- | --- | --- | --- | --- |
|  | pH 5.5 | pH 7.5 | pH 5.5 | pH 7.5 |  |
| 17-257 | + | + | + | - | + |
| 17-259 | + | + | + | - | - |
| 19-060 | + | + | + | - | - |
| 19-061 | + | + | + | - | - |
| 19-063 | + | + | + | - | - |

### 3.2. Identification of C. auris polypeptides secreted at pH 5.5 and pH 7.5 using high-throughput mass-spectroscopy

To identify polypeptides secreted by *C. auris* at pH 5.5 (SECR5) and pH 7.5 (SECR7), *C. auris* cells were grown in minimal media containing amino acids and glucose as a carbon source. The pH of each medium was maintained by adding 100 mM MES (pH 5.5) or HEPES (pH 7.5). Proteins present in cell-free fractions of three biological replicates were independently concentrated, subjected to trypsin digestion and further analysed by mass-spectroscopy (MS) according to the workflow depicted in Fig. S1. Further processing of MS data (for details, see Materials and Methods) revealed 123 *C. auris* proteins possessing signal polypeptides that were considered as members of the putative *C. auris* secretome (Table S1). In addition to proteins exclusively present in SECR5 (13 in total) or SECR7 (27 in total), 83 proteins were common, although their abundance varied between SECR5 and SECR7. Further analysis of their abundance by VolcaNoseR (https://huygens.science.uva.nl/VolcaNoseR2/) made it possible to identify those that are preferentially enriched in SECR5 and SECR7 (Fig. 1).

**Fig. 1.**
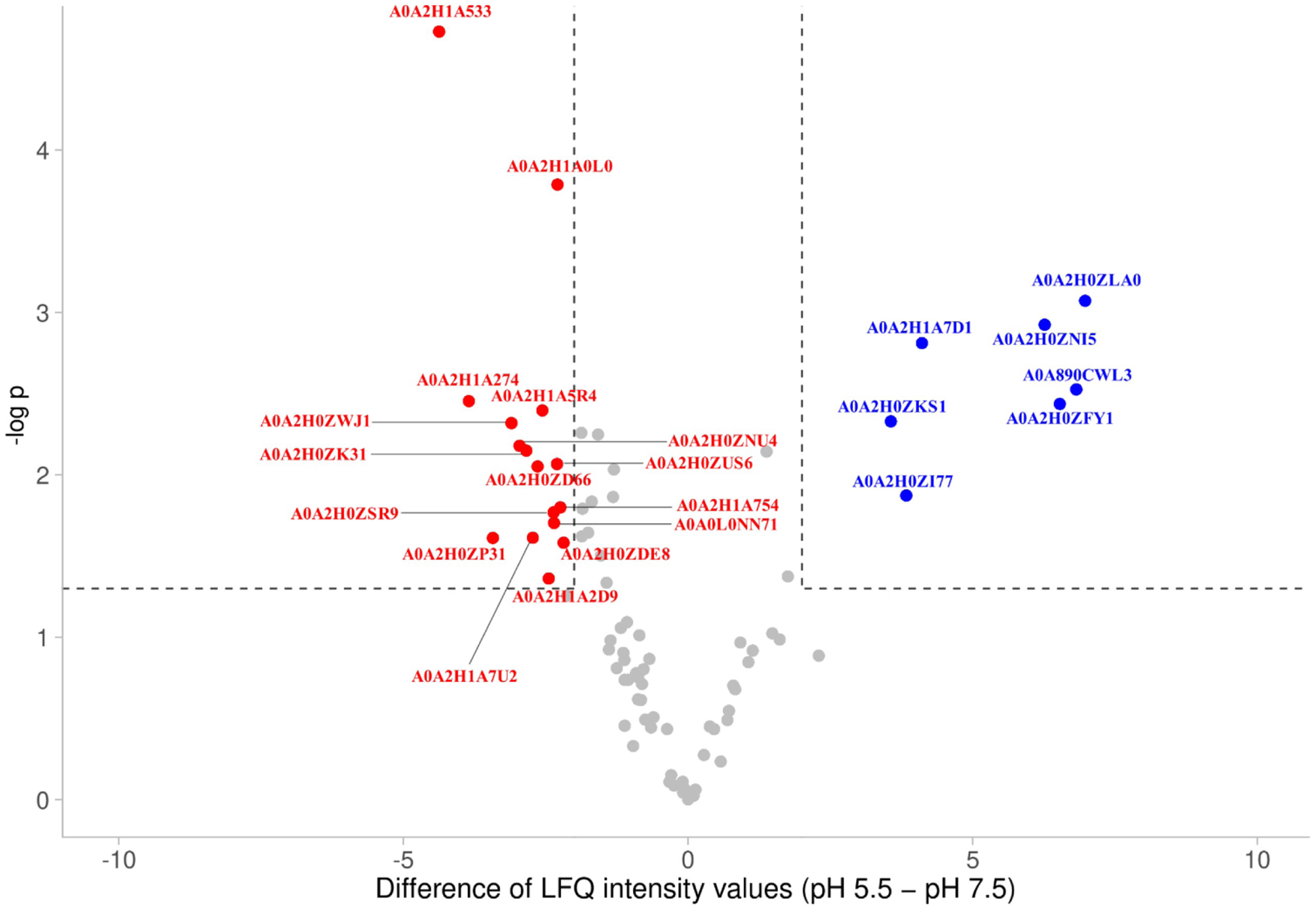
Volcano plot of 83 secreted proteins possessing signal polypeptides. Secreted proteins (see Table S1) that were significantly enriched (p < 0.05) in SECR5 and SECR7 are indicated by closed red and blue circles, respectively. Statistical analysis was carried out by using a two-tailed paired t-test.

### 3.3. Gene ontology analysis of C. auris secretome

Gene ontology (GO) analysis of 123 secreted proteins representing the combined secretome (i.e., both SECR5 and SECR7) was carried out using CGD Gene Ontology Term Finder (http://www.candidagenome.org/cgi-bin/GO/goTermFinder). This analysis revealed that the secreted proteins of *C. auris* could be annotated within a number of GO terms (Table S2). The largest group (28 proteins) represent polypeptides involved in fungal-type cell wall organization / biogenesis (GOID 71852 and 71554) followed by polypeptides with known functions in carbohydrate metabolism (GOID 5975). Comparison of *C. auris* secretome (this study) with that of *C. albicans* [35] revealed that they have 37 proteins in common (Fig. 2A). This finding suggests that a large fraction of the secretome appears to be evolutionarily conserved among these pathogens. Furthermore, GO analysis using the aforementioned Gene Ontology Term Finder revealed that the common polypeptides are represented by various cell wall-modifying enzymes as well as those with proteolytic activities (Fig. 2B). Nevertheless, *C. auris* secretome contains a large fraction of unique proteins (86 in total) including those potentially involved in modulation of host immune response, virulence and cell wall biogenesis.

**Fig. 2.**
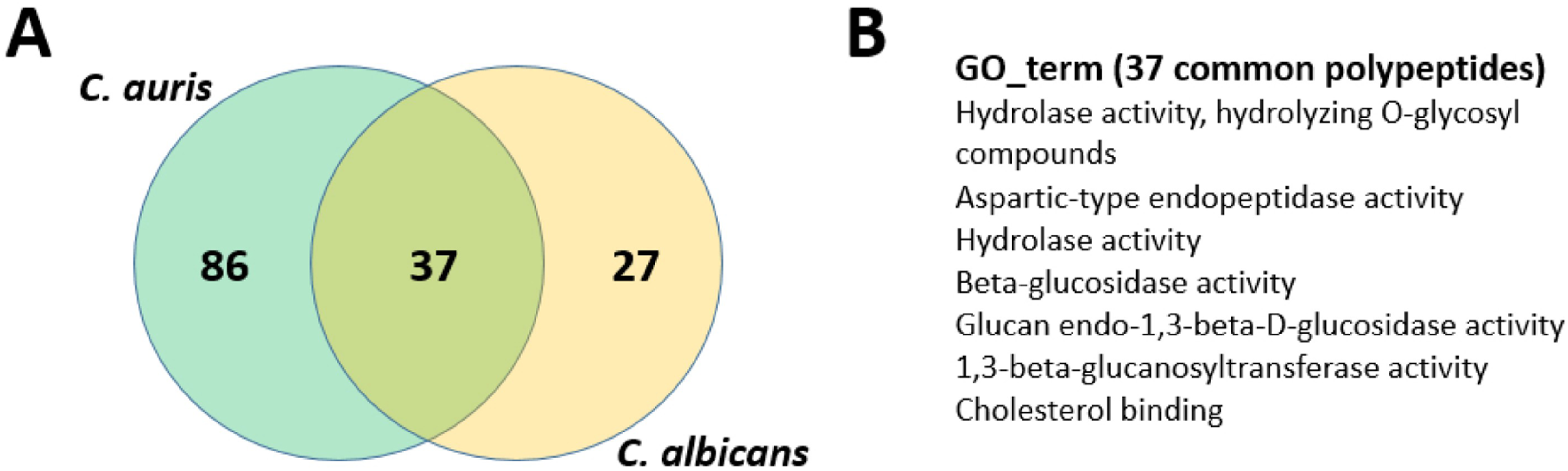
Comparative analysis of secreted *C. auris* and *C. albicans* polypeptides. **(A)** The Venn diagram shows the overlap in the secreted proteins of *C. auris* and *C. albicans*. **(B)** GO terms that describe the secreted polypeptides common for the two species.

### 3.4. Defining protein-interacting networks

Bioinformatics analysis of the combined secretome (i.e., *C. auris* proteins present in cell-free fractions at both pH 5.5 and pH 7.5) using STRING package [36] yielded a long network of polypeptides; small protein clusters and a considerable number of standalone proteins. Dissection of the long protein network into smaller clusters (Fig. 3) revealed that the corresponding proteins are likely involved in several important processes related to cell wall biogenesis and host-pathogen interactions.

**Fig. 3.**
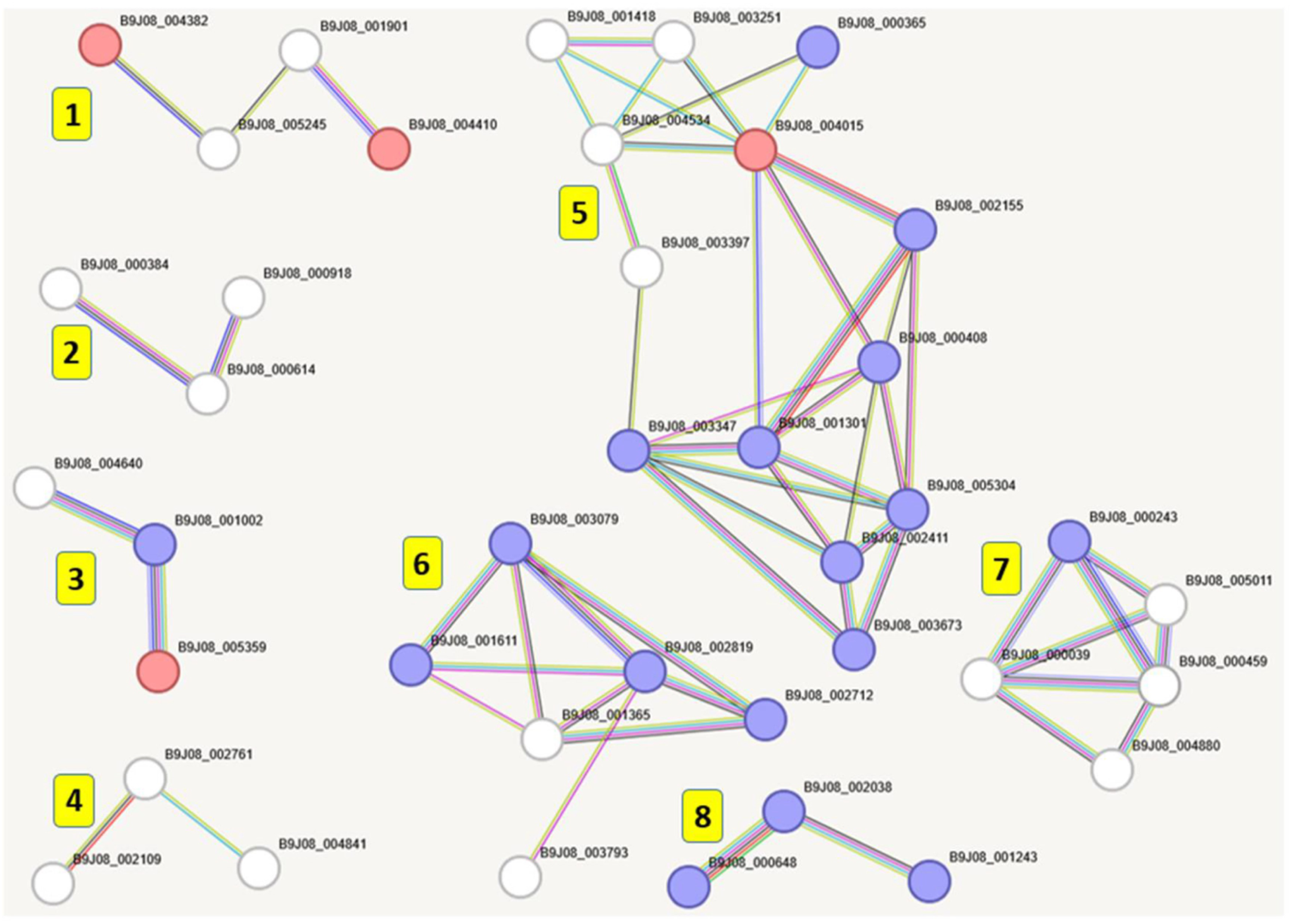
Functional clusters identified within the main protein network of *C. auris* secretome by using K-mean clustering. The original protein network deduced by STRING and its splitting into 8 functional clusters using K-mean clustering. The depicted protein clusters combine proteins (i) hydrolyzing O-glycosyl compounds (cluster 1), (ii) processing polypeptides in endoplasmic reticulum (cluster 2), (iii) possessing unknown activities (cluster 3), (iv) impacting fungal-type cell wall (cluster 4), (v) involved in amino sugar and nucleotide metabolism (cluster 5) and (vi) representing *C. auris* rotamase (cluster 6), receptor L-domain superfamily (cluster 7) as well as those controlling mixed pathways (cluster 8). Proteins that present only in SECR5, SECR7 or shared by both SECR5 and SECR7 are indicated by red (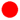), blue (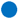), or white (0) circles, respectively.

### 3.5. Differential secretion of polypeptides at pH 5.5 versus pH 7.5

Although many polypeptides found in the secretomes obtained at both pH conditions, i.e., in SECR5 and SECR7, were the same and likely represent the core of the *C. auris* secretome, some proteins (Table S1) were exclusively present (or significantly enriched) in SECR5 (13 polypeptides), whereas others in SECR7 (27 polypeptides). The differences in protein content of SECR5 and SECR7 have a significant impact on the composition of two major clusters (i.e., clusters 5 and 6 shown Fig. 3). Namely, we found that the more “complete” variants of these clusters were seen only for polypeptides present in SECR7 (Fig. 3). Moreover, some enzymes, for example, dolichol phosphate mannose synthase (i.e., enzyme composed of three subunits (B9J08_005304, B9J08_002411 and B9J08_003673) and ancillary factors (e.g., glycoprotein glucosyl transferase Gpt1, B9J08_000408) are present only in SECR7 (indicated by blue circles in cluster 5, Fig. 3). These enzymes play an essential role in protein N- and O-glycosylation [37]. Similarly, components of the disulphide isomerase (B9J08_002712), peptidyl-prolyl isomerase (B9J08_002038 and B9J08_000648) and heat-shock protein 70 (Hsp70, B9J08_001611) that are part of cluster 6 (Fig. 3) were present in SECR7 but not in SECR5. These above proteins function in protein folding, and they were likewise identified in *C. albicans* secretome [38].

Unlike SECR7, SECR5 contains a number of polypeptides that were only present (or were significantly enriched) at pH 5.5 (Fig. 4). They were arbitrarily placed into four groups including (i) aspartic proteases (e.g., Sap2, Sap3 and so on), (ii) putative virulence factors (e.g., agglutinine-like protein), (iii) several cell-wall proteins as well as (iv) uncharacterized peptides with unknown functions (Fig. 4).

**Fig. 4.**
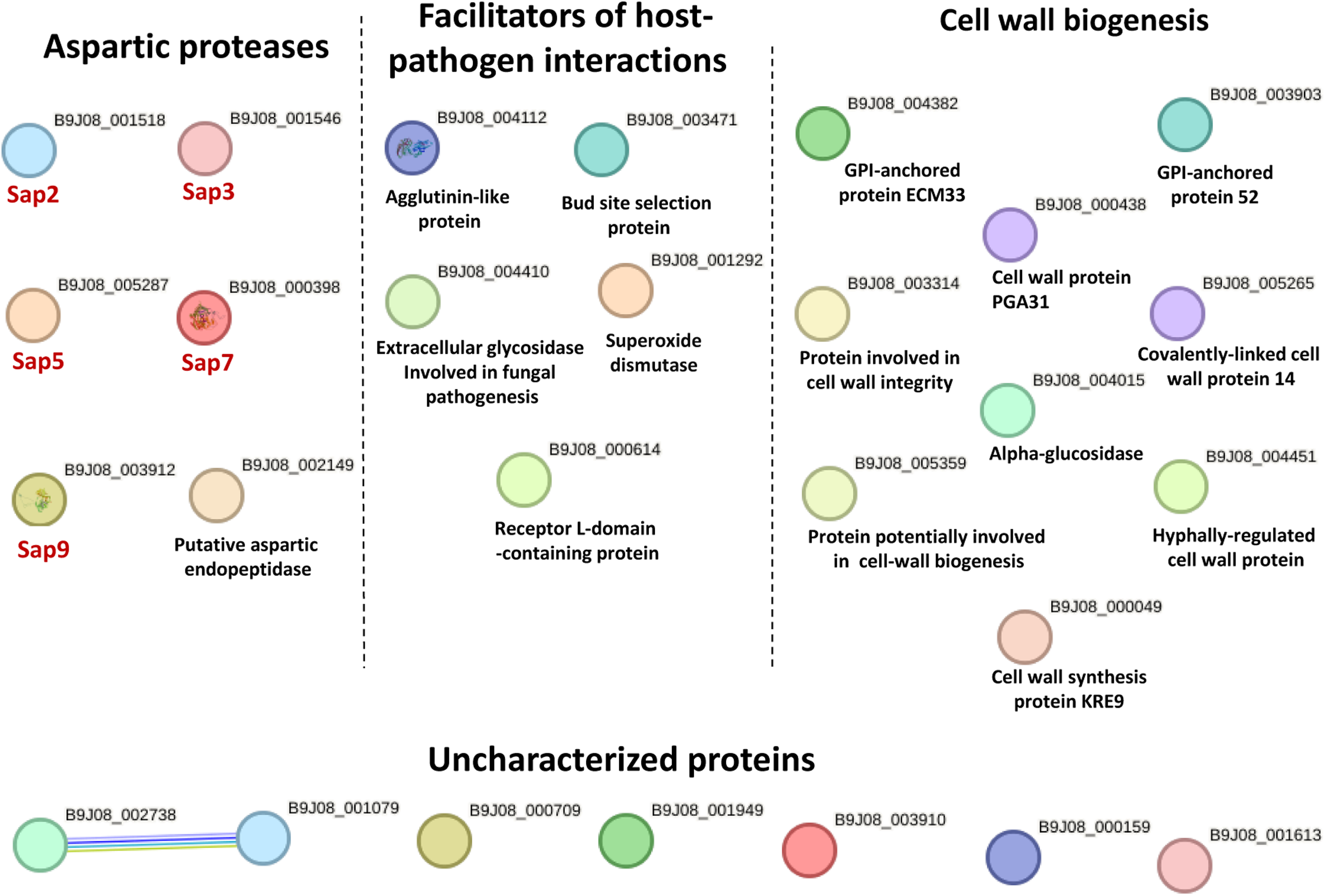
Secreted *C. auris* polypeptide that are exclusively present or significantly enriched in SECR5. These proteins include aspartic proteases, different facilitators of host-pathogen interactions, cell wall proteins as well as 7 poorly annotated polypeptides.

## 4. Discussion

The significant contribution of *Candida-*related species such as *C. auris* to nosocomial diseases raised great interest in studying host-pathogen interactions involving these fungi [5]. Previous studies revealed a number of secreted virulence factors potentially playing important roles in *Candida*-associated diseases [13,39]. Although recent analyses of *Candida* secretomes enabled to provide deeper understanding of virulence factors, the comprehensive identification and analysis of virulence factors, whose secretion is mediated by signal peptides, have been performed only for a few *Candida* species. Moreover, two recent studies of *C. auris* were primarily focused on the composition and biological functions of the extracellular vesicles containing a large variety of macromolecules (i.e. numerous metabolic enzymes [40] and RNA molecules [41]) trapped in the cytoplasm. Although the above studies shed some light on the biological roles of extracellular vesicles, still little is known about secretion of the major *C. auris* hydrolytic enzymes (i.e., proteases, lipases and haemolysins) and other putative virulence factors secreted on their own (i.e., secreted independent of vesicles due to the presence of signal peptides) known for their essential roles in *Candida*-associated diseases.

Pilot plate assays revealed that some *C. auris* clinical isolates of our strain collection secret polypeptides possessing protease- and lipase-like activities (Table 1). To learn more about the nature of the corresponding enzymes and disclose other protein factors secreted by this pathogen, we decided to analyse the composition of *C. auris* secretome using a clinical isolate (i.e., 17-257), which belongs to clade III of this species. Moreover, we carried out analysis of two different secretomes obtained for *C. auris* cells grown in media with pH 5.5 and pH 7.5, respectively, thus mimicking different pH conditions potentially faced by this pathogen in the host. Mass-spectroscopy identification of polypeptides released by *C. auris* cells into minimal medium at pH 5.5 and pH 7.5 revealed 123 proteins possessing signal peptides. Similar to production of extracellular proteins by *C. albicans* [38] and *Nakaseomyces glabratus* (previously known as *Candida glabrata* [29,42]), *C. auris* secretes polypeptides known for their role in cell wall integrity, maintenance and biogenesis (Table S1). These polypeptides along with other secreted putative virulence factors might play a crucial role in colonization of hosts and pathogen survival. Indeed, we found that a number of secreted polypeptides represent hydrolytic enzymes potentially facilitating host tissue damage, nutrient acquisition and pathogen survival in the host. Namely, the presence of various glucanases, lipases, phospholipases and aspartic proteinases in the analysed secretomes (see Table S1) suggests that *C. auris* can process host-derived macromolecules (e.g., extracellular polysaccharides, polypeptides and lipids) by breaking them into smaller molecules potentially utilized by this pathogen as carbon and nitrogen sources. This scenario is supported by recent studies demonstrating that disruption of genes coding for aspartic proteases [43] or lipases [44] can negatively affect *Candida* virulence and/or survival in the host environment. Besides their essential roles in nutrient acquisition, some hydrolytic polypeptides along with a number of cell wall modifying enzymes can contribute to cell wall morphogenesis [22,23], thus playing an essential role in infection and pathogen persistence in the host [24]. Unlike the cell wall of other fungal pathogens, *C. auris* cell wall possesses some unique properties that let this pathogen efficiently evade its recognition by the host immune system [10]. It is conceivable that the evasion may include cell wall modification at all three layers including mannan, glucan and chitin. In agreement with this idea, we found that the *C. auris* secretome contains the putative enzymes able to control biosynthesis of mannan (e.g., various mannan endo-1,6-alpha-mannosidases (B9J08_003852 and B9J08_002188) and manosyl-oligosaccharide glucosidase (B9J08_003347)), chitin (e.g., chitinase (B9J08_002761) and chitin deacetylase (B9J08_002761)) as well as β-1,3- and β-1,6-glucans (e.g., exo-beta-1,3-glucanase (B9J08_003799) and 1,3-beta-glucanosyltransferase (B9J08_005245)).

The high plasticity of the cell wall and secretome enables *C. auris* to adapt to different environments and host-pathogen interactions that occur in various anatomical sites differing in nutrient content and pH [45]. We demonstrate here that pH affects the composition of SECR5 and SECR7, both containing a number of unique polypeptides. This observation is consistent with previous findings demonstrating that pH can affect the composition and activity of secretomes produced by *C. albicans* and other *Candida* species (see [46] and references therein). Analysis of *C. auris* secretome revealed that protein clusters 5 and 6 (Fig. 3) were particularly enriched in SECR7-specific polypeptides. For example, some SECR7-specific polypeptides in cluster 5 include those that represent three subunits of dolichol phosphate mannose synthase (i.e., B9J08_005304, B9J08_002411 and B9J08_003673), which contributes to protein glycosylation and *Candida* morphogenesis [37]. The other protein cluster (cluster 6) contains SECR7-specific polypeptides with the putative role in protein folding.

In contrast, a number of proteins including aspartic proteases were exclusively found in SECR5 (Fig. 6). The secretion of aspartic proteases at pH 5.5 is consistent with the previous observations indicating that *Candida* species secret this type of proteases at acid pH. Noteworthy, one of them, Sap3, was previously defined as a primary aspartic protease that greatly contributes to *C. auris* virulence and is involved in biofilm formation [15]. Other SECR5-specific virulence factors likely play an essential role in host-pathogen interactions. For example, the agglutinin-like protein (B9J08_004112) might facilitate attachment of the pathogen to a host-tissue prior to biofilm formation, whereas the bud-site selection protein (B9J08_003471) can stimulate haploid-invasive growth [47].

The pH-sensing and rewiring of gene expression in *Candida* spp. is regulated by the Rim101-dependent signal transduction pathway [48]. Previous work has shown that the activation of Rim101 pathway can enhance expression of chitinases in *C. albicans* [49], thereby altering the recognition of this pathogen by the innate immune system. We found that an ortholog of *C. albicans* chitinase (i.e., B9J08_002761) is present in the *C. auris* core secretome. Further work is necessary to clarify whether this enzyme plays the same role in *C. auris* cell wall remodelling as it does in *C. albicans*.

During yeast-like cell growth, cell wall undergoes tremendous changes leading to a release of some wall-bound proteins into the medium. Consequently, the detached proteins such as GPI proteins can be detected in supernatant obtained after cell pelleting [50,51]. Although their detachment is sometimes associated with mechanical alteration in the cell wall structure, the presence of GPI proteins in cell-free fractions can also be attributed to other factors, for example, it can be caused by the action of the aspartic proteases Sap9 and Sap10 [52]. We found that the number of GPI proteins was increased from 4 to 7 in SECR5, i.e., when *C. auris* was grown in more acid environment. Moreover, the concomitant finding of 4 extra aspartic proteases (AP) in SECR5 raises the possibility that detachment of new GPI proteins in SECR5 might be due to their partial proteolysis by APs.

## 5. Conclusion

Taken together, the above findings suggest that *C. auris* conditionally produces a number of secreted virulence factors that might differentially affect the capacity of this pathogen to colonize host surfaces (skin and mucosae) and elicit infection in a pH-dependent manner.

## CrediT authorship contribution statement

**Asier Ramos-Pardo:** Methodology, Formal analysis and investigation, Writing - original draft preparation. **Guillermo Quindós:** Conceptualization, Writing - review and editing, Funding acquisition. **Elena Eraso:** Conceptualization, Writing - review and editing, Funding acquisition. **Elena Sevillano:** Conceptualization, Methodology, Formal analysis and investigation, Writing - original draft preparation, Writing - review and editing, Funding acquisition. **Vladimir Kaberdin:** Conceptualization, Methodology, Formal analysis and investigation, Writing - original draft preparation, Writing - review and editing, Funding acquisition. All authors commented on previous versions of the manuscript. All authors read and approved the final manuscript.

## Data availability statement

All raw data have deposited in the ProteomeXchange Consortium via the PRIDE partner repository with the dataset identifier PXD060071 and doi 10.6019/PXD060071. The raw data can be accessed by logging into the PRIDE website (https://www.ebi.ac.uk/pride/login) using the following details: Project accession number: PXD060071 Token: 7L90VVHi1j2c

## Acknowledgements

The authors thank Dr. Kerman Aloria for assisting with the mass-spec analysis of polypeptides performed in the Proteomics Core Facility-SGIKER at the University of the Basque Country (member of ProteoRed-ISCIII). The authors also thank Javier Pemán and Alba Ruiz-Gaitán (Hospital Universitario y Politécnico La Fe, Valencia, Spain) for kindly providing clinical *C. auris* isolates.

