## Supplementary material for "Effect of pH on the secretome profile of the human pathogen *Candidozyma auris*": Figure S1

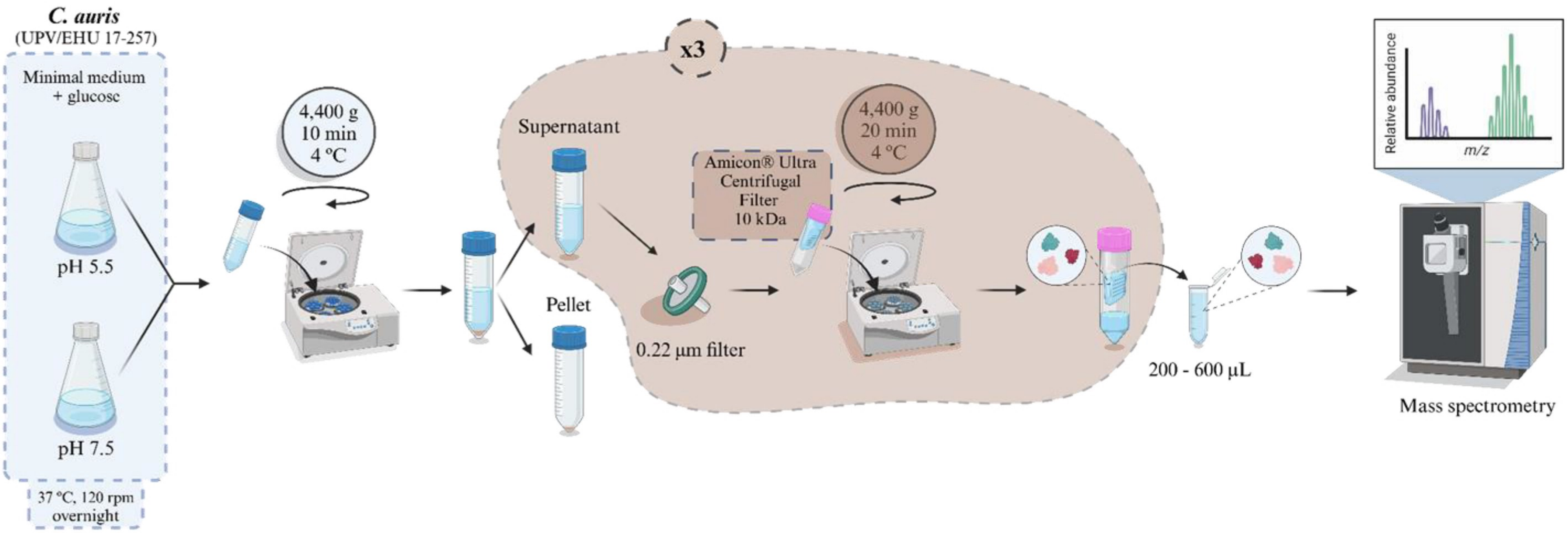


**Fig. S1. Preparation of protein samples for mass-spectroscopic analysis of polypeptides secreted by *C. auris*.** The image was created in https:/BioRender.com.
